# The implied motion aftereffect changes decisions, but not confidence

**DOI:** 10.1101/498048

**Authors:** Regan M. Gallagher, Thomas Suddendorf, Derek H. Arnold

## Abstract

Viewing static images depicting movement can result in a motion aftereffect: people tend to categorise direction signals as moving in the opposite direction to the implied motion of still photographs. This finding could indicate that inferred motion direction can penetrate sensory processing and impact perception. Equally possible, however, is that inferred motion impacts decision processes, but not perception. Here we test these two possibilities. Since both categorical decisions and subjective confidence are informed by sensory information, confidence can be informative about whether an aftereffect likely results from changes to perceptual or decision processes. We therefore leveraged subjective confidence as an additional measure of the implied motion aftereffect. In Experiment 1 (implied motion), we find support for decision-level changes only, with no change in subjective confidence. In Experiment 2 (real motion), we find equal changes to decisions and confidence. Our results suggest the implied motion aftereffect produces a bias in decision-making, but leaves perceptual processing unchanged.

---

An outstanding question in perception research is whether our thoughts, desires, emotions, or cognitions can change how our sensory systems operate. Usually, perceptual aftereffects are quantified by recording subjective responses to a range of stimulus intensities (e.g. the brightness of lights, or tone frequencies), and measuring decision changes before and after prolonged and repeated exposures to a specific stimulus (see Clifford, Webster, Stanley, Stocker, Kohn, & Sharpee et al., 2007 for a review). Aftereffects are also typically constrained to a common sensory dimension, such as when a moving adaptor influences the perceived motion of a test (Barlow & Hill, 1963). However, recent research has started revealing aftereffects in which test stimuli are only *conceptually* related to the adapting stimulus. These studies offer an empirical approach to determine whether high-level cognitions (that extract meaning) can change perception.

One study conducted by Winawer, Huk & Boroditsky (2008) had participants adapt to still photographs that implied motion, either leftward or rightward, or inward or outward. In a second study (Winawer, Huk & Boroditsky, 2010), participants adapted to a static grating stimulus, and were asked to imagine that it was moving. In both studies, participants then judged the motion direction of a dynamic dot stimulus. In each case, the adaptation phase (static images, or imagined motion) gave rise to a *negative* aftereffect: participants were more likely to judge a (possibly ambiguous) stimulus as having moved in the opposite direction. This pattern of results is broadly consistent with the classic motion aftereffect (Barlow & Hill, 1963), and the authors concluded that the adaptation task had directly impacted motion perception. While this interpretation is intuitive, there is an equally plausible interpretation: viewing static images that imply movement, or imagining movement, might engage motion-related cognitions that bias categorical decisions, but leave the sensory processes underlying motion perception unchanged.

Recent research suggests that subjective confidence reports can provide an important additional source of information to help discern whether or not an aftereffect is likely to have a perceptual basis (Gallagher, Suddendorf, & Arnold, 2018). When decisions are impacted by sensory adaptation, both the decision and confidence functions can provide equivalent measures of the perceptual aftereffect. However, when decisions change for reasons other than a perceptual change (e.g. due to a biased pattern of response when inputs are ambiguous), the range of inputs that elicit low confidence in categorical judgments can remain veridical, and a dissociation between decision and confidence response profiles can emerge. In this scenario, the dissociation constitutes evidence for the aftereffect being driven by decisional processes.

Subjective confidence can be helpful for diagnosing the origin of an aftereffect by providing a second source of information that is directly informed by sensory evidence. Moreover, confidence can be dissociated from categorical decisions when decision biases change categorisation responses (see Gallagher et al., 2018). Measuring changes in confidence, in addition to category responses, can thus help to distinguish changes in perception from changes in decision-making. In the present study, we find evidence that viewing still photographs depicting movement changes categorisations of ambiguous inputs, but not confidence. On this basis, we argue that that the implied motion aftereffect is unlikely to be a perceptual aftereffect.

## Perceiving versus deciding

Biases in decision-making are usually accepted as the evidence for sensory adaptation, but these can equally occur in the absence of perceptual changes (see Morgan et al., 2011). Based on biased decisions alone, we cannot tell whether a given aftereffect has resulted from changes to perception or from changes to a decision process independent of perception (for greater discussion on this topic, see Morgan et al, 2011; Storrs, 2015; Firestone & Scholl, 2016; Fritsche et al., 2017). To illustrate, Figure 1 depicts two components of perceptual decision-making: sensory evidence and decisional criteria.

**Figure 1:**
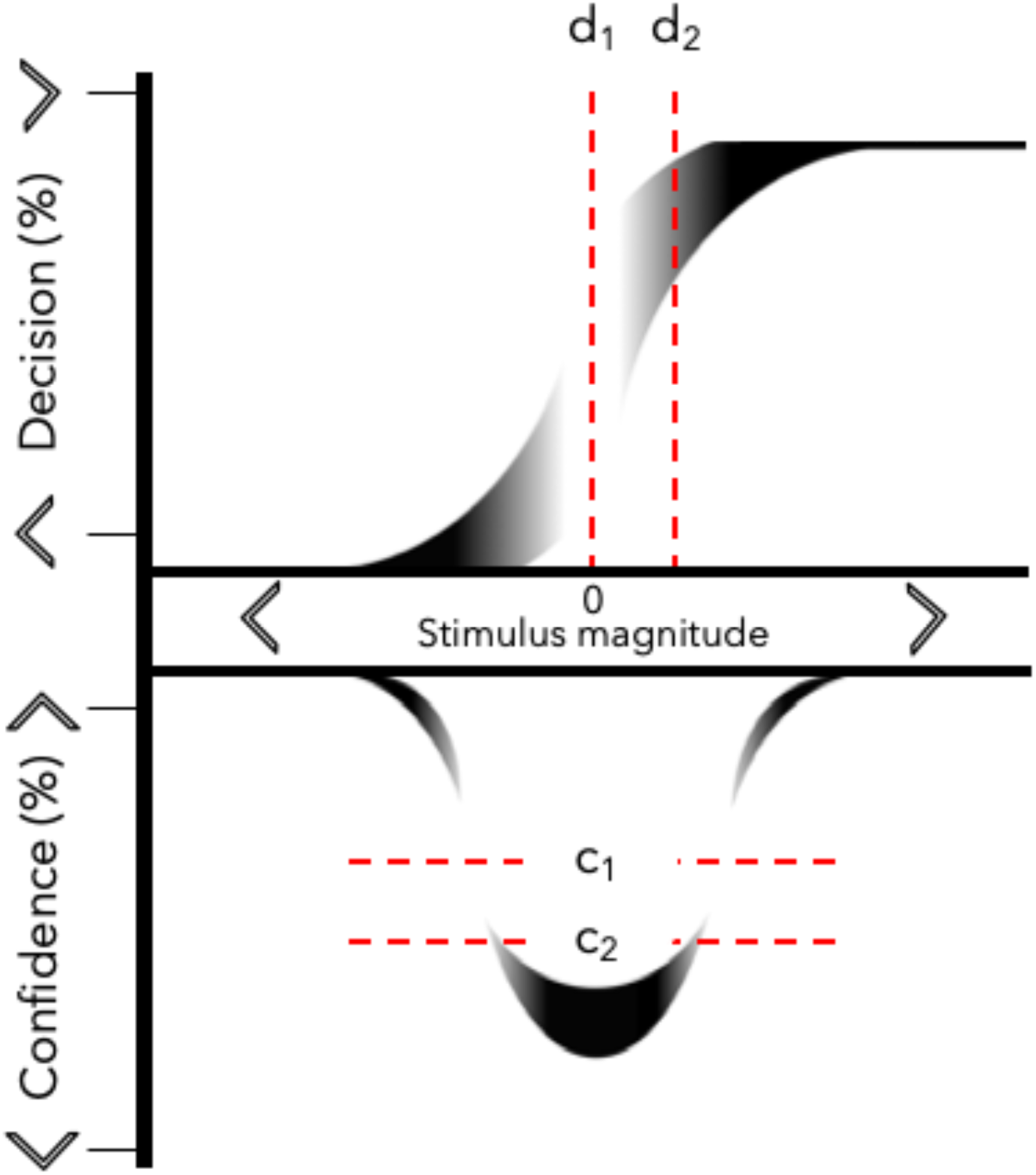
A discrimination task with two binary responses: decisions, left or right, and confidence, high or low. (*top*) A sigmoid function (S-curve) represents the sensory signal-to-noise ratio (with greater noise implied by lighter shading) plus response criteria (red dotted lines). As the stimulus intensity approaches an encoded value of 0, the observer’s decisions become more probabilistic. The placement of decision criteria impacts central tendency estimates for the decision distribution (d_1_ = unbiased, d_2_ = left-response bias). (*bottom*) A Gaussian function (U-curve) representing confidence in decision-making. As the stimulus intensity approaches 0, confidence decreases to its minimum. Response criteria for confidence relate to the strength of evidence needed to report confidence, which can encourage more or less reports of confidence, but do not encourage a lateral shift of the confidence function (c_1_ = low-confidence threshold, c_2_ = high-confidence threshold).

Sensory evidence consists of the physiological information made available to the brain by the sense organs. Sensory evidence is typically considered to be contaminated by Gaussian noise, so different sensory information can be encoded on a trial-by-trial basis, even if people are repeatedly exposed to an unchanging physical input. Decision criteria can be regarded as a boundary, or threshold, separating when people will decide whether to classify a stimulus as belonging to one category or another (as moving to the left or right in this context). The interpretation of weaker stimuli (with encoded values close to a decision boundary) can be shifted to either side of the criterion on a trial-by-trial basis by sensory noise, by a changing decision criterion, or both. So, decisions made about weak signals are often probabilistic. Stronger signals might be equally variable from trial-to-trial, but variable decision or sensory processes will have less impact on their interpretation, because the encoded value is more distant from the criterion.

If an observer is required to choose between two alternatives when an input is perceptually ambiguous, they may rely on higher-order cognitive (i.e. non-perceptual) aspects of decision-making. One might simply guess, leading to an equal likelihood of choosing left or right. Alternatively, they might adopt some other strategy that carries greater bias toward one or another category. A decision strategy could manifest as either a negative or a positive bias (or none), independent of sensory evidence and other post-perceptual processes (see Yarrow et al, 2011). Moreover, a wilfully adopted decision bias can change estimates of decision boundaries, without impacting on the precision of perceptual judgments (Morgan et al., 2011) or the central tendency of the associated distribution of confidence (Gallagher et al., 2018).

## The Present study

A participant whose task is to imagine or infer movement while viewing stationary objects might, when prompted, show a statistical preference for the unimagined direction when test stimuli are ambiguous (Winawer et al., 2008; Winawer et al., 2010). In this example, the observer could be indexing unchanged sensory evidence against a different criterion when uncertain. Crucially, humans can report on whether they are confident or guessing, which can provide important additional information about how to interpret otherwise-ambiguous changes in categorical responses.

Changes in decision-making could occur without physiological changes in motion-sensitive brain regions, due to a shift in decision criteria. This would alter the measurement of maximal categorical ambiguity—the inflection point of a cumulative gaussian function fit to binary categorical decisions. This measure is often referred to as the point of subjective equality (PSE), and changes in this metric from baseline trials are used to estimate aftereffect magnitudes. However, while a shift in a decision criterion will alter decision-making, they might have no impact on the range of inputs that elicit uncertainty. An additional important point is that both measures (decisions and confidence) are informed by sensory encoding. So, having people report on decisional confidence, in addition to committing to a categorisation, might reveal a dissociation. Confidence, or expressions of uncertainty, could remain veridical in the case where categorisations are biased solely by decisional processes.

In two Experiments we measure categorical direction decisions and confidence reports to help determine whether the implied motion aftereffect likely results from changes to perceptual or post-perceptual decision processes.

## Aim and hypotheses

The aim of the current study is to test whether implied motion from still images changes perception of real motion. We use confidence measures to provide additional diagnostic information. In Experiment 1, we measure motion-direction decisions following adaptation to a stream of still photographs depicting directional motion (either to the left or right). In Experiment 2, we compare this implied motion aftereffect to a more conventional motion aftereffect, using a serial dependence between successive real motion signals. We hypothesise that changes to sensory processing will impact categorical direction decisions and confidence reports equally. In contrast, we hypothesise that aftereffects arising from post-perceptual (decision) processes will *selectively* impact direction decisions—non-perceptual changes in decision-making should produce a dissociation between decision and confidence responses.

## Experiment 1

### Method

#### Participants

All participants (30 for each Experiment) were recruited from the University of Queensland’s Psychology department. Sample sizes of 30 were set for all Experiments, as these are comparable to samples used in the original studies of the imagined and implied motion aftereffects. Participants were drawn from a first-year student pool, who received course credit for their participation. All were naïve as to the purpose of the experiments.

#### Ethics

Ethical approval for all experiments was obtained from the University of Queensland’s Ethics Committee, and experiments were conducted in accordance with committee guidelines. Each participant provided written informed consent, and were aware that they could withdraw from the study without penalty at any point.

#### Materials & Stimuli

Stimuli were presented on a Dell LCD monitor (1024×768 pixels). All computers were running Matlab software and the Psychophysics Toolbox (Brainard, 1997; Pelli, 1997). All monitors had a screen refresh rate of 60Hz. Adapting stimuli (Experiment 1 only) and test stimuli were presented within an aperture size of 300 × 175 pixels against a grey background (RGB = 125,125,125).

Adapting stimuli were photographs taken from a Google Image search, after searching for images that depicted fast movement along the horizontal plane. The implied motion stimuli were then compiled into a set of 100 photographs, with an approximately equal aspect ratio in portrait orientation. The dimensions of photographs were re-sized before presentation, so that they were all equal in aspect ratio. Each photo was mirror-flipped, so it could be used to depict both directions.

Test stimuli consisted of 100 dots rendered blue against a grey background. Each dot was 1 pixel in size. Initially, dots were drawn at random locations within the aperture window. Dot coherence values ranged from −30 (30% coherence leftward) through 0 (random motion) to +30 (30% coherence rightward). Test stimuli were set to one of 11 coherence values [−30 −20 −10 −6 −3 0 3 6 10 20 30], presented in a randomised order. Coherently moving dots were selected at random on each frame, so no individual dot could be tracked across the screen. Coherent motion was achieved by displacing coherent dots left or right by one pixel on successive frames. All other dots were redrawn at random locations.

#### Procedure

Participants sat comfortably in a chair approximately 55 cm from the display, resting their hands on the keyboard’s directional buttons while fixating a central cross-hair. If there was an adaptation phase, the adapting stimulus was presented for 18 seconds on the first trial for each of five blocks, and again on the middle trial when the implied motion direction reversed. For all other trials the adapting stimulus lasted for six seconds. Participants passively viewed the implied motion images without responding. See Figure 3 for a representation of the task procedure.

After the adaptation phase, participants were presented with one of the 11 dot-motion test probes. Tests were presented for one second before disappearing, leaving only the fixation cue. Participants reported whether the test had appeared to be moving left (by pressing the left arrow key) or right (by pressing the right arrow key). If participants could not determine the test direction, they were instructed to make their best guess.

Once the direction judgment had been made on each trial, the fixation cue turned black, prompting a confidence response. Participants indicated whether they felt high confidence in their direction judgment by pressing the up arrow on the keyboard, or low confidence (or that they were guessing) by pressing the down arrow. The fixation cross turned white once the confidence response had been provided, and a new trial started immediately. Each of the 11 stimulus values was tested five times per block, and participants completed two blocks in total (once with a leftward adaptor and once with a rightward adaptor) resulting in 110 individual test trials per participant. The initial adaptor direction was randomised for each participant.

**Figure 2.**
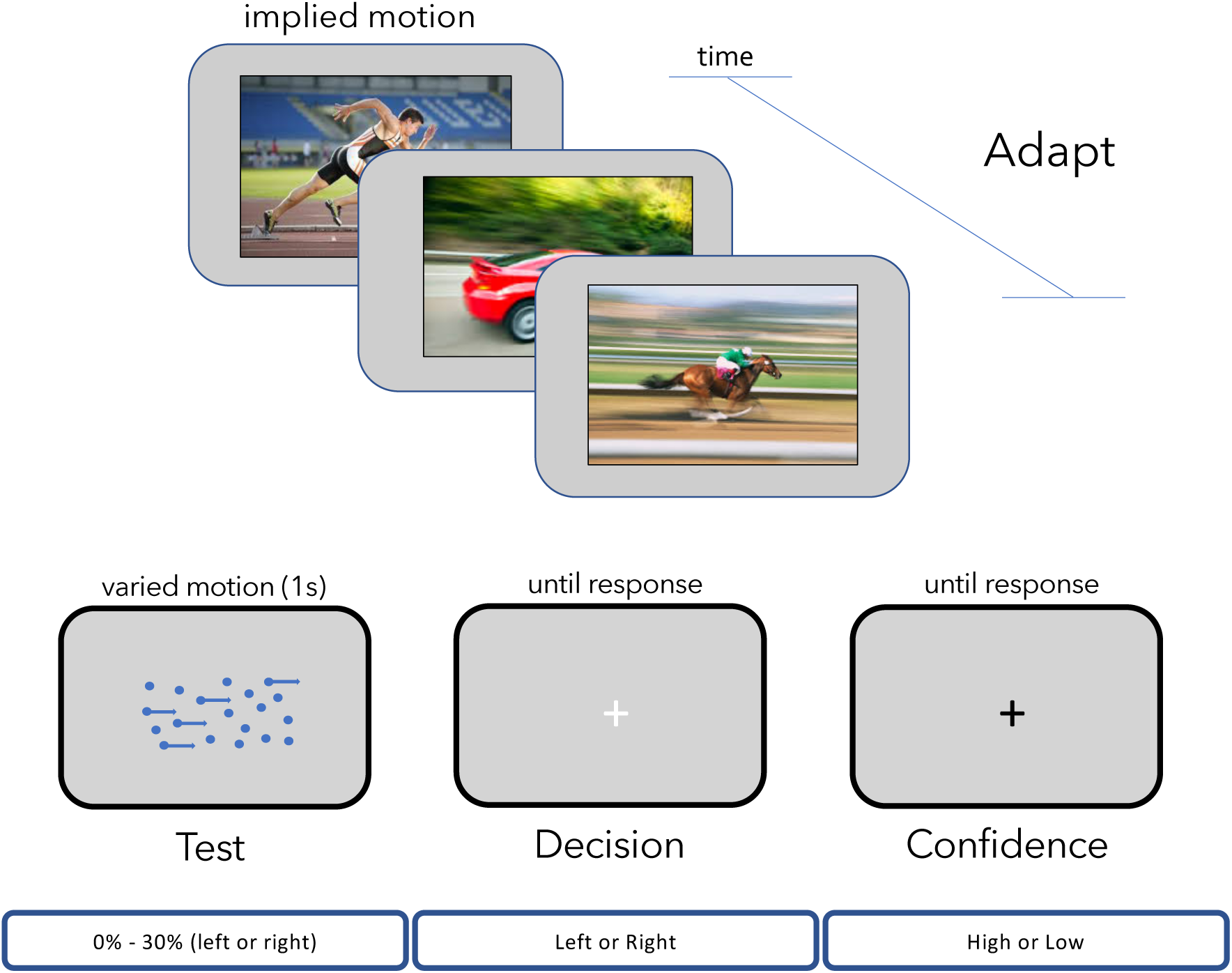
Experiment procedure. In Experiment 1, participants adapted to still photographs that depicted motion to either the left or right (one direction for each testing block). Adapting stimuli appeared for 18s on the first trial of each block, and on the middle trial, and for 6s on all other trials. There was no adaptation period in Experiment 2. Each trial then had a dot test probe. Tests were present for 1s, appearing on the second frame after the adapting stimulus disappeared. A new trial began once participants had recorded their direction decision (left or right) and reported their confidence in their decision (high or low).

#### Data preparation

Categorical direction decisions were scored as a 1 if the person said the stimulus moved to the right, or 0 if they said left. The response function approximated a Sigmoid distribution when the proportion of rightward decisions is plotted as a function of motion direction and coherence. The inflection point of the Sigmoid function (for an unbiased observer) should approximate 0% coherence, indicating an equal probability of choosing leftward and rightward motion when the stimulus has random physical movements.

Confidence responses were scored as a 0 if the person reported being confident in their direction decision, or as −1 if they were not confident. This response distribution approximated a raised Gaussian function (bell curve) when responses were plotted as a function of motion direction and coherence. The peak of the confidence function is the point of *lowest* confidence, which should be centred around 0% coherence for an unbiased observer. Two aftereffect measures were taken for each participant in each Experiment: one from the inflection point of decision responses, and one from the peak of confidence responses.

### Results

All *t*-tests reported below are two-way repeated-measures tests for equality of means. All Bayes Factor analyses were estimated using JASP software (JASP, 2016) with a Cauchy prior width of 0.707.

#### Implied motion aftereffect

Analyses for Experiment 1 showed that adaptation to still images depicting motion had a robust impact on direction decisions. Results showed that decisions following leftward adapting images (L_PSE_ = 0.71; SD = 2.70) were significantly different from decisions following rightward adapting images (R_PSE_ = 2.50; SD = 4.70; *t*_29_ = 2.94, *p* = .006, Cohen’s d = 0.54, 95% CI 0.15 – 0.91, BF_10_ = 6.56). However, implied motion adaptation did not impact measures of confidence. The central tendency of confidence reports following leftward implied motion adaptors (L_CONF_ = 0.84; SD = 1.50) was not significantly different to the central tendency of confidence reports following rightward implied motion adaptors (R_CONF_ = 0.78; SD = 2.29; *t*_29_ = 0.13, *p* = .896, BF_10_ = 0.20). We also measured a dissociation between the effect of implied motion on decision (∆PSE = 1.79; SD = 3.33) and confidence reports (∆CONF = −0.06; SD = 2.55; *t*_29_ = 2.50, *p* = .018, Cohen’s d = 0.46, 95% CI 0.08 – 0.83, BF_10_ = 2.71). These data are depicted in Figure 3.

**Figure 3:**
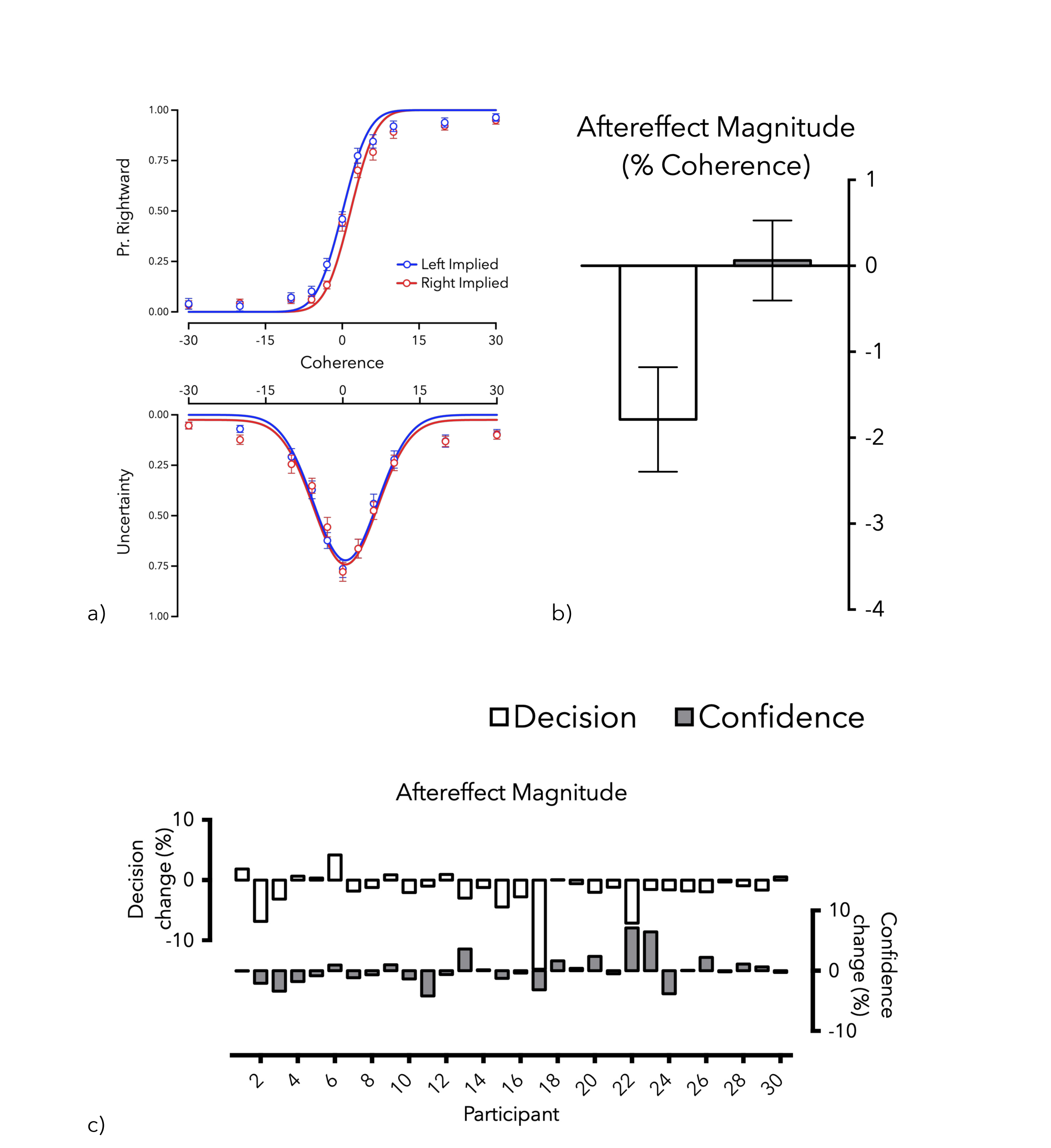
Results of Experiment 1. (a) psychometric functions depicting (*top*) the proportion of rightward responses and (*bottom*) the proportion of confidence responses, both as a function of motion direction and coherence. Separate lines depict direction of the implied motion adaptor. (b) average aftereffect magnitudes, as measured by decision and confidence. (c) Individual data showing changes in decision and confidence following implied motion adaptation. N = 30; error bars depict ± 1 s.e.m.

## Experiment 2

### Method

The methodological details for Experiment 2 are identical to Experiment 1 except for the following. One, participants sequentially responded to the test stimuli without an adapting stimulus. Two, there were eight test values [−30 −15 −5 −1 +1 +5 +15 +30] rather than the 11 test values in Experiment 1. There was no 0% coherence value, which allowed us to analyse responses according to the physical direction of the previous test. Three, participants completed 50 observations of each of the 8 test values for a total of 400 observations per observer.

### Results

#### Rapid motion aftereffect

Analyses for Experiment 2 showed that the test motion direction on the previous trial had a robust impact on subsequent direction decisions. Results showed that decisions following leftward tests (L_PSE_ = −1.02; SD = 2.85) were significantly different from decisions following rightward tests (R_PSE_ = 1.05; SD = 2.31; *t*_27_ = 3.41, p = .002, Cohen’s d = 0.65, 95% CI 0.23 – 1.05, BF_10_ = 18.09). The previous test motion direction also impacted measures of confidence. The central tendency of confidence reports following leftward tests (L_CONF_ = −0.95; SD = 2.23) was significantly different to the central tendency of confidence reports following rightward tests (R_CONF_ = 1.76; SD = 2.92; *t*_27_ = 3.38, *p* = .002, Cohen’s d = 0.64, 95% CI 0.23 – 1.04, BF_10_ = 16.77). The measured effect of previous trials was not significantly different for decision (∆PSE = 2.07; SD = 3.21) and confidence reports (∆CONF = 2.71; SD = 4.24; *t*_27_ = 0.70, *p* = .488, BF_10_ = 0.25). These data are depicted in Figure 4.

**Figure 4.**
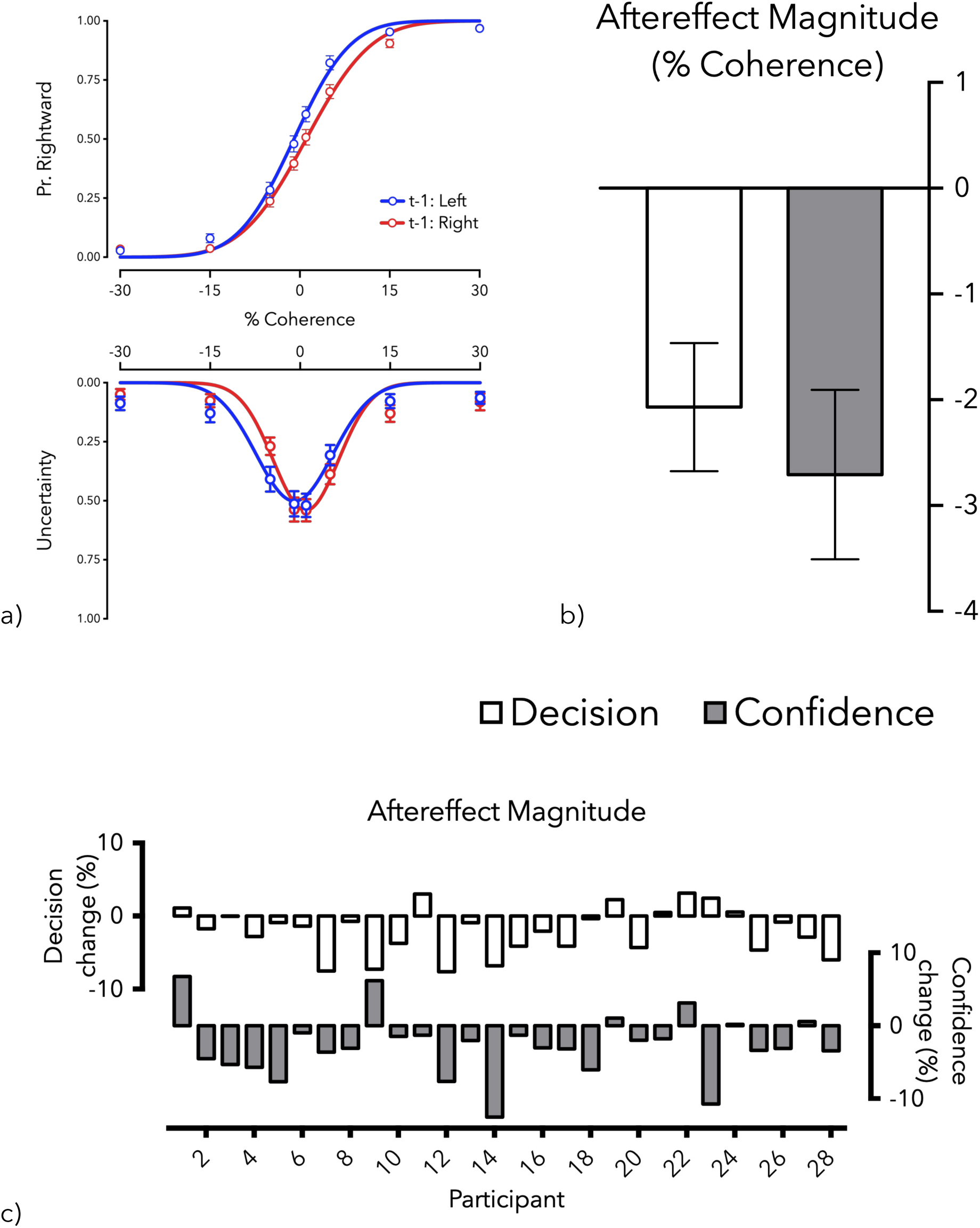
Results of Experiment 2. (a) psychometric functions depicting, top, the proportion of rightward responses and, bottom, proportion of confidence responses as a function of motion coherence. Separate lines depict direction of the previous motion stimulus. (b) average aftereffect magnitude as measured by decision and confidence. (c) Individual data showing changes in decision and confidence following rapid motion adaptation. N = 28; error bars depict ± 1 s.e.m.

## Discussion

Our results suggest that the implied motion aftereffect is the result of changes to post-perceptual decision processes. Experiment 1 replicated the effect of viewing a stream of still photographs implying directional motion. As predicted, and consistent with previous research (Winawer et al., 2008), our results showed that participants were more likely to report that tests were moving in the *opposite* direction relative to that implied by still adapting images. The results of confidence responses, however, suggested that the region of peak uncertainty was unchanged by adapting to implied motion, and a dissociation between direction decisions and confidence reports was observed.

Experiment 2 showed that the PSE and confidence measures were *equally* influenced by the previous physical test motion. Tests on the current trial (trial *n*) were more likely to be reported as moving in the *opposite* direction relative to the last test (trial *n*-1). The probability that the current test stimulus would evoke low confidence was also impacted by the previous test stimulus. Importantly, the shift in confidence reports was in the *same* direction as the shift in decisions, and of equal magnitude. These results are consistent with both responses (direction decisions and confidence) being informed by a common source of information (perception), which was equally impacted by recent sensory history.

Previous research shows that perceptual changes result in an equivalent impact on the central tendency of both decision and confidence reports (Gallagher et al., 2018), whereas changes in post-perceptual processes can change decision-making without changing confidence. The pattern of results in Experiment 2 is consistent with evidence for a perceptual aftereffect, whereas the decision aftereffect in Experiment 1 is consistent with changes in non-perceptual processes.

Our results show that the degree to which implied motion (Experiment 1) and serial dependence (Experiment 2) impact motion direction decisions was approximately equal (both Cohen’s d effect sizes ~0.6). However, only exposure to real motion produced a change in confidence reports. This finding is important in at least two ways. First, it shows that confidence measures are sufficiently sensitive to detect moderate sized changes in decision-making, ruling out the possibility that confidence is not sensitive enough to detect perceptual changes from viewing still photographs. Second, it shows that aftereffects that elicit equal changes in categorical decision-making (compare the decision aftereffect estimates of Exp1 and Exp2) can nonetheless be separated based on how they impact another response (confidence) which also relies on sensory evidence. These results point to the utility of confidence in the study of perceptual aftereffects.

A core finding in our serial dependence results is in agreement with previous research showing that perceived motion can undergo rapid sensory adaptation (Kanai & Verstraten 2005; Glasser et al., 2011), and that rapid negative motion aftereffects have an equivalent impact on the central tendency of both decision and confidence reports (Gallagher et al., 2018). While our serial dependence results showed a negative aftereffect, many other serial dependencies have revealed positive aftereffects (Fischer & Whitney; Cicchini, Mikellidou, & Burr; Fornaciai & Park, 2018), which are thought to produce a more stable representation of our dynamic world, although it remains a matter of debate whether this stability is a decision-level effect rather than a perceptual experience (Fritsche et al., 2017). Judgements of confidence could help reveal whether other serial dependencies likely have a perceptual basis.

Aftereffects induced by implying or imagining motion have previously been interpreted as resulting from a top-down impact of cognition on motion perception (Winawer et al., 2008, 2010; Pavan et al., 2011). This claim is bolstered by electrophysiological evidence, that viewing photographs depicting implied motion activates some of the same direction-selective cortical circuits as viewing real motion (Lorteije et al., 2006; Lorteije et al., 2007). However, it is not clear from either set of evidence that motion perception has been changed by cognition. Instead, different decisions could be reached, and similar neural activations could be observed, by changes to decision processes, even in the absence of changes to motion perception.

Although our data cannot determine why implied motion adaptation produces a systematic negative aftereffect, previous research demonstrates that arbitrary decision strategies can produce data consistent with either a positive or a negative aftereffect (Morgan et al., 2011; Gallagher et al., 2018). One possibility is that people surreptitiously perform a categorisation task whenever an unambiguous signal is encountered. This could result in a bias toward categorising ambiguous inputs in the *opposite* category, relative to the more recently viewed (adapting) direction signal. This describes the frequency principle—a propensity to assign equal numbers of inputs to either category when making dichotomous classifications (Parducci et al., 1960; Parducci & Wedell, 1986). So, if a given class of input is repeatedly ‘implied’ as an adapting signal, people might compensate by assigning more weight to the less frequent category.

## Conclusion

Our data suggest that the implied motion aftereffect reflects changes in decision-making, independently of changes to sensory processes. This contrasts with the motion aftereffect that changes both directional decisions and estimates of directional uncertainty.

## Authorship declaration

RMG and DHA designed the experiments. RMG conducted the experiments, analysed the data, and wrote the manuscript. RMG, TS, and DHA revised and approved the final manuscript. The authors declare no conflict of interest.

## References

Barlow, H. B., & Hill, R. M. (1963). Evidence for a physiological explanation of the waterfall phenomenon and figural after-effects. Nature, 200(4913), 1345.

Barlow, H. B., & Hill, R. M. (1963). Selective sensitivity to direction of movement in ganglion cells of the rabbit retina. Science, 412–414.

Brainard, D. H., & Vision, S. (1997). The psychophysics toolbox. Spatial vision, 10, 433–436.

Clifford, C. W., Webster, M. A., Stanley, G. B., Stocker, A. A., Kohn, A., Sharpee, T. O., & Schwartz, O. (2007). Visual adaptation: Neural, psychological and computational aspects. Vision research, 47(25), 3125–3131.

Firestone, C., & Scholl, B. J. (2016). Cognition does not affect perception: Evaluating the evidence for" top-down" effects. Behavioral and brain sciences, 39.

Fritsche, M., Mostert, P., & de Lange, F. P. (2017). Opposite effects of recent history on perception and decision. Current Biology, 27(4), 590–595.

Gallagher, R. M., Suddendorf, T., & Arnold, D. H. (2018). Confidence as a diagnostic tool for perceptual aftereffects. bioRxiv, 270280.

JASP Team. (2016). JASP (Version 0.7. 5.5) [Computer software]. Google Scholar, 765, 766.

Lorteije, J. A., Kenemans, J. L., Jellema, T., Van Der Lubbe, R. H., De Heer, F., & Van Wezel, R. J. (2006). Delayed response to animate implied motion in human motion processing areas. Journal of Cognitive Neuroscience, 18(2), 158–168.

Lorteije, J. A., Kenemans, J. L., Jellema, T., Van der Lubbe, R. H., Lommers, M. W., & van Wezel, R. J. (2007). Adaptation to real motion reveals direction-selective interactions between real and implied motion processing. Journal of Cognitive Neuroscience, 19(8), 1231–1240.

Mollon, J. (1974). After-effects and the brain. New Scientist, 61(886), 479–482.

Morgan, M., Dillenburger, B., Raphael, S., & Solomon, J. A. (2012). Observers can voluntarily shift their psychometric functions without losing sensitivity. Attention, Perception, & Psychophysics, 74(1), 185–193.

Parducci, A.,Calfee, R.C., Marshall, L.M. & Davidson, L.P. (1960). Context effects in judgment: Adaptation level as a function of the mean, mid-point, and median of the stimuli. Journal of Experimental Psychology 60, 65 – 77.

Parducci, A. & Wedell, D.H. (1986). The category effect with rating scales: Number of categories, number of stimuli, and method of presentation. Journal of Experimental Psychology: Human Perception and Performance 12, 496 – 516.

Pavan, A., Cuturi, L. F., Maniglia, M., Casco, C., & Campana, G. (2011). Implied motion from static photographs influences the perceived position of stationary objects. Vision research, 51(1), 187–194.

Pelli, D. G. (1997). The VideoToolbox software for visual psychophysics: Transforming numbers into movies. Spatial vision, 10(4), 437–442.

Prism, G. (2017). version 7.0 e. GraphPad Software.

Storrs, K. R. (2015). Are high-level aftereffects perceptual?. Frontiers in psychology, 6, 157.

Tanner Jr, W. P., & Swets, J. A. (1954). A decision-making theory of visual detection. Psychological review, 61(6), 401.

Winawer, J., Huk, A. C., & Boroditsky, L. (2008). A motion aftereffect from still photographs depicting motion. Psychological Science, 19(3), 276–283.

Winawer, J., Huk, A. C., & Boroditsky, L. (2010). A motion aftereffect from visual imagery of motion. Cognition, 114(2), 276–284.

Yarrow, K., Jahn, N., Durant, S., & Arnold, D. H. (2011). Shifts of criteria or neural timing? The assumptions underlying timing perception studies. Consciousness and cognition, 20(4), 1518–1531.

